# NED-Net: open-source software for building and refining seizure detectors during rodent EEG experiments

**DOI:** 10.64898/2026.10.15.759362

**Authors:** Mohd Yaqub Mir, Marco Ledri

## Abstract

Long-term EEG is the main readout of rodent epilepsy models, and automated detectors are usually delivered as fixed models that a laboratory cannot adapt to its own animals. NED-Net is open-source software in which annotations, rule-based detections and model proposals share one file per recording, reviewed events train the next detector, and detection runs live during acquisition, so a detector can be refined while an experiment runs. A detector built in NED-Net on one kainate mouse cohort detected 64% of 221 video-adjudicated convulsive seizures in a second cohort as delivered, and 72% of its detections on held-out recordings were seizures.

Retraining on annotations made in the app, a few seconds of review per event, raised recall to 88% at 74% precision. SeizyML, a published detector, reached similar recall with three times the detections and did not improve with the same annotations. A convulsive classifier removes non-seizures and enriches convulsive events.

## Introduction

Chronic video-EEG monitoring is a major outcome measure in rodent models of acquired epilepsy, and the volume of recorded data has outgrown what humans can review manually. The two experiments from which the cohorts analyzed in the present study were drawn together comprise almost 3,000 recording files and over 30,000 animal-hours of EEG.

Several automated detectors for rodent EEG have been published and perform well on their development datasets. A classifier over 22 engineered features reached sensitivity and positive predictive value above 98% in three mouse models^1^, and a generalized linear model over 141 features reached an area under the receiver operating characteristic curve (AUROC, the probability that a randomly chosen seizure window is scored higher than a randomly chosen non-seizure window) of 0.995 on 1,012 spontaneous seizures in rats^2^. The latter study also performed external validation on an independently constructed cohort of 96 rats, where the AUROC fell to 0.890. A classifier developed on three mouse models retained 91–99% sensitivity within those models but 76% on a fourth model it had not been trained on^3^, and in a rat model of post-traumatic epilepsy a deep convolutional network trained on each animal individually outperformed the same architecture trained across animals (seizure-detection accuracy, the fraction of visually scored seizures detected: 0.960 versus 0.811)^4^. Taken together, these studies indicate that detector performance can decline on recordings outside the development set and may vary substantially between animals.

The practical question for a laboratory is therefore not whether a published detector is accurate on its development data, but how to obtain a detector that works on its own recordings, how to check it, and how to improve it when it falls short, preferably while the experiment is still running. Tools with a graphical interface exist for rodent seizure detection and review^1,5^, and SeizyML in particular pairs interpretable classifiers trained on the user’s own labels with manual validation of every detected event^5^. In these tools, the validated events are the endpoint. To our knowledge, none return them to training, attribute them to animals so that review can be directed at the implants where the detector is weak, or run during acquisition.

Here we describe NED-Net (Neural Event Detection Network), open-source software built around an annotation–training–review loop, and we evaluate it on two mouse cohorts recorded in our laboratory. We ask three questions. Does a detector built in the software on one cohort transfer to a new cohort as delivered? What does running the loop on the new cohort cost in annotation, and what does it add? And how does the result compare with a published tool run on the same recordings against the same reference? The software is not seizure-specific, in the sense that it can be used to train a network to detect theoretically any EEG signal of interest.

Additionally, it does not necessarily require machine learning. It ships also with rule-based detectors, and its review interface and per-animal reporting form a complete workflow on their own.

## Results

### NED-Net closes the loop between annotation and detection

NED-Net is built around a single data structure, a sidecar annotation file stored next to each EDF recording, in which human annotations, rule-based detections, and model proposals share one format (Fig. 1, interface in Fig. 2). A laboratory without a trained model begins with rule- based candidate detection: the detectors propose candidate events on the first recordings, the reviewer confirms, rejects, or adds events in a review queue, and the resulting annotations train the first U-Net. The model’s detections return to the same queue as pending events, and reviewing them produces the next training set, so the second and subsequent models are trained the same way as the first. Because each channel is mapped to an animal, every event is attributed to an animal from the first file, and all summaries are reported per animal rather than per file. Detection can process completed recordings in a folder, on a cluster, or live as the acquisition system writes files. Per-animal detector output is therefore visible within days of implantation, while there is still time to annotate the animals that need it and retrain before most of the protocol has been recorded.

**Figure 1.**
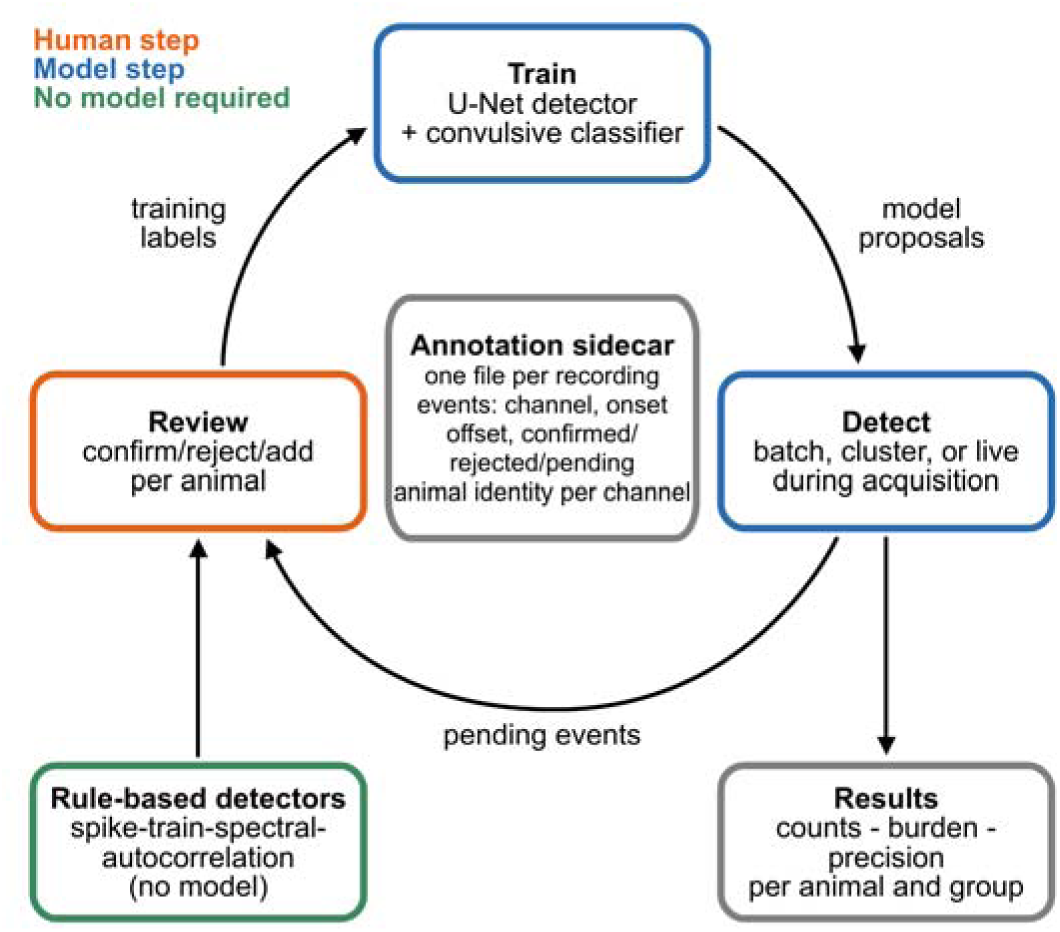
The annotation–training loop implemented by NED-Net. NED-Net stores all human annotations, rule-based detections, and model proposals in a single annotation sidecar per recording (center), which records each event’s channel, onset, offset, and label (confirmed, rejected, or pending), along with each channel’s animal identity. A laboratory without a trained model starts with the rule-based detectors (dashed green, no model required), whose proposals are reviewed per animal (orange, human step). Confirmed and rejected events become the training labels of a U-Net detector (Stage 1) and a convulsive classifier (Stage 2) (blue, model step). The trained cascade runs on completed recordings in batch, on a computing cluster, or live during acquisition, and its proposals return to the same review queue as pending events, so that each round of review produces the training set of the next model. Per-animal counts, burden and precision are aggregated in the Results view.

**Figure 2.**
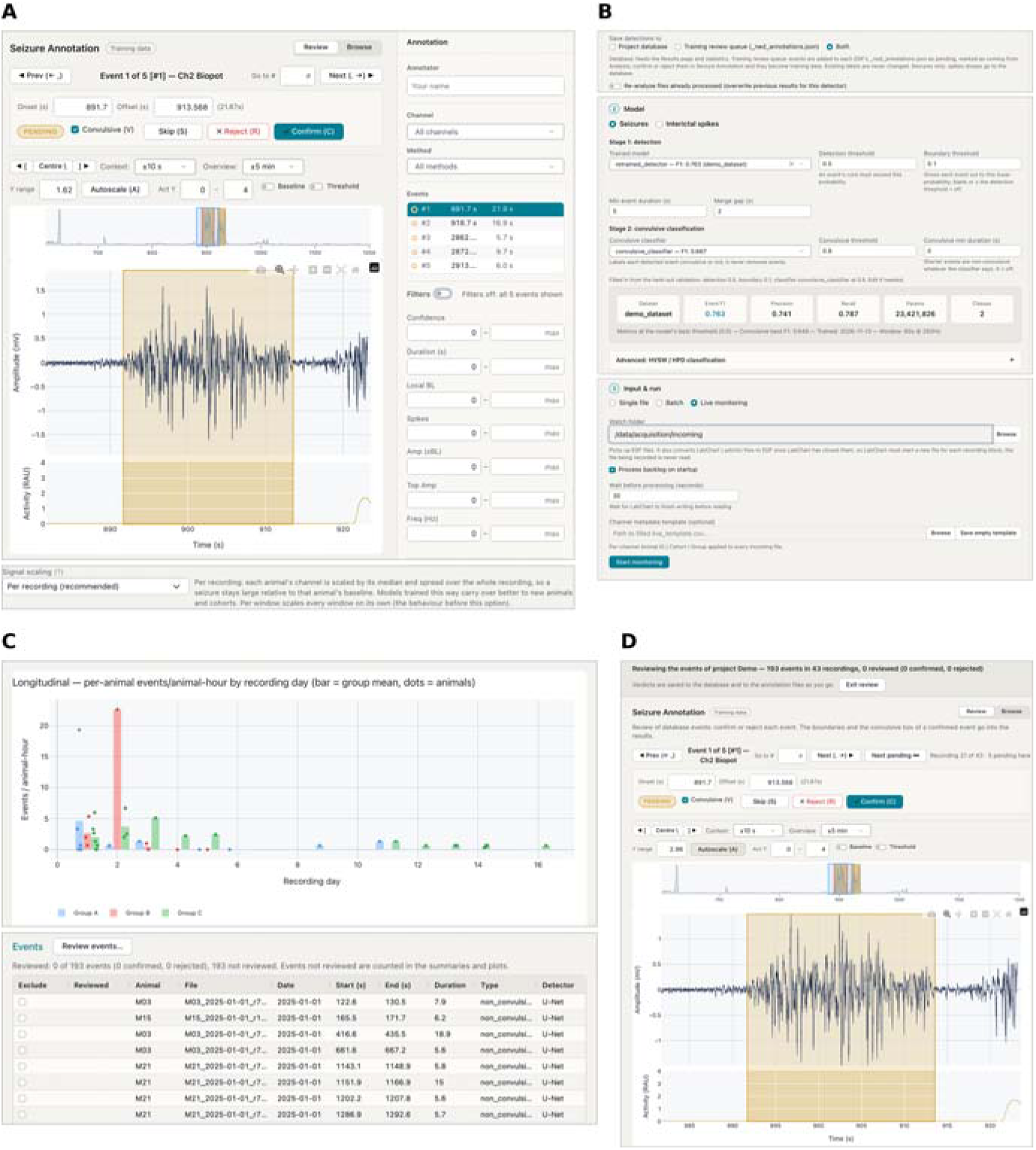
The NED-Net interface. (A) Annotation and review in the Training tab: a pending U- Net detection in a second-cohort recording (animal M05, 21.9 s, detector confidence 0.84) is shown with the EEG trace, the activity channel, the detected extent (shaded), an overview of the surrounding minutes, and the confirm, reject and skip controls, with a checkbox that marks the event convulsive. Boundaries can be dragged on the trace and manual events added. The strip below shows the signal scaling option of the training form, per recording by default. (B) The Analysis tab: detections are written to the project database, to the annotation sidecars for review, or to both. A trained detector and convulsive classifier are selected together with their operating point (detection threshold, boundary threshold, minimum duration and merge gap), filled in from the validation of the selected model, and the mode selector switches between single file, batch and live monitoring of a watched folder with backlog processing. (C) The Results tab: events per animal-hour by recording day with per-animal points and group means, one of several views (Supplementary Fig. 1), and the event list of the run with the control that starts the review of its detections. (D) Review of a run’s detections from the Results tab, in the same view as (A). Each detection is confirmed, rejected or skipped, rejected detections are removed from the counts, and verdicts are written to the project database and to the sidecars, where they can feed the next training round.

We used this loop on two cohorts of kainate-treated mice recorded on the same acquisition system. A detector was trained on the first cohort and applied unchanged to the second, the second cohort was then annotated through the interface, and the detector was retrained. A reviewer blinded to NED-Net output adjudicated candidate events in the second cohort on synchronized video, identifying 221 convulsive seizures on the retained channels (21 animals, 7,142 recording-animal units, 10,461 h), which served as the reference for recall. Precision was measured by review of every detection made on 55 recording-animal units that the retrained model had not been trained on (Methods). Throughout, a reference seizure counts as detected when a detection overlaps it within 5 s, both detectors run at the same operating point (detection threshold 0.5, boundary threshold 0.1), and operating points were chosen as the threshold with the highest event-level F1 on each model’s validation data.

## A detector built on one cohort transfers to the next

The source detector was trained from annotations made in the interface on the first cohort (867 confirmed seizures from 165 recordings), with the EEG of each recording normalized by a center and scale computed once per channel over the whole recording (Methods). On its validation split, it reached an event-level F1 of 0.83 at threshold 0.5 (precision 0.75, recall 0.93), and on three source-cohort animals whose seizures were not used in training, it detected 380 of 408 confirmed seizures (93%), and 96% of its 689 detections were seizures (95% interval 93–97%, Methods).

The same detector, at the same operating point, was then run on the second cohort (Table 1). It detected 141 of the 221 convulsive reference seizures (64%), producing 5,544 detections in 10,461 recording-hours (0.53 per hour) and flagging 17.6 h as seizure. Among the 11 animals with at least five reference seizures, per-animal recall ranged from 42% to 91% with a median of 65%, and no animal was left without a detected seizure (Fig. 3, Supplementary Table 2). On the 55 held-out recording-animal units, which hold 30 reference seizures, it detected 19, and 90 of its 125 detections there were seizures on review (72%). Recall was unchanged without the 5 s tolerance and one seizure lower under one-to-one matching (Supplementary Table 1).

**Figure 3.**
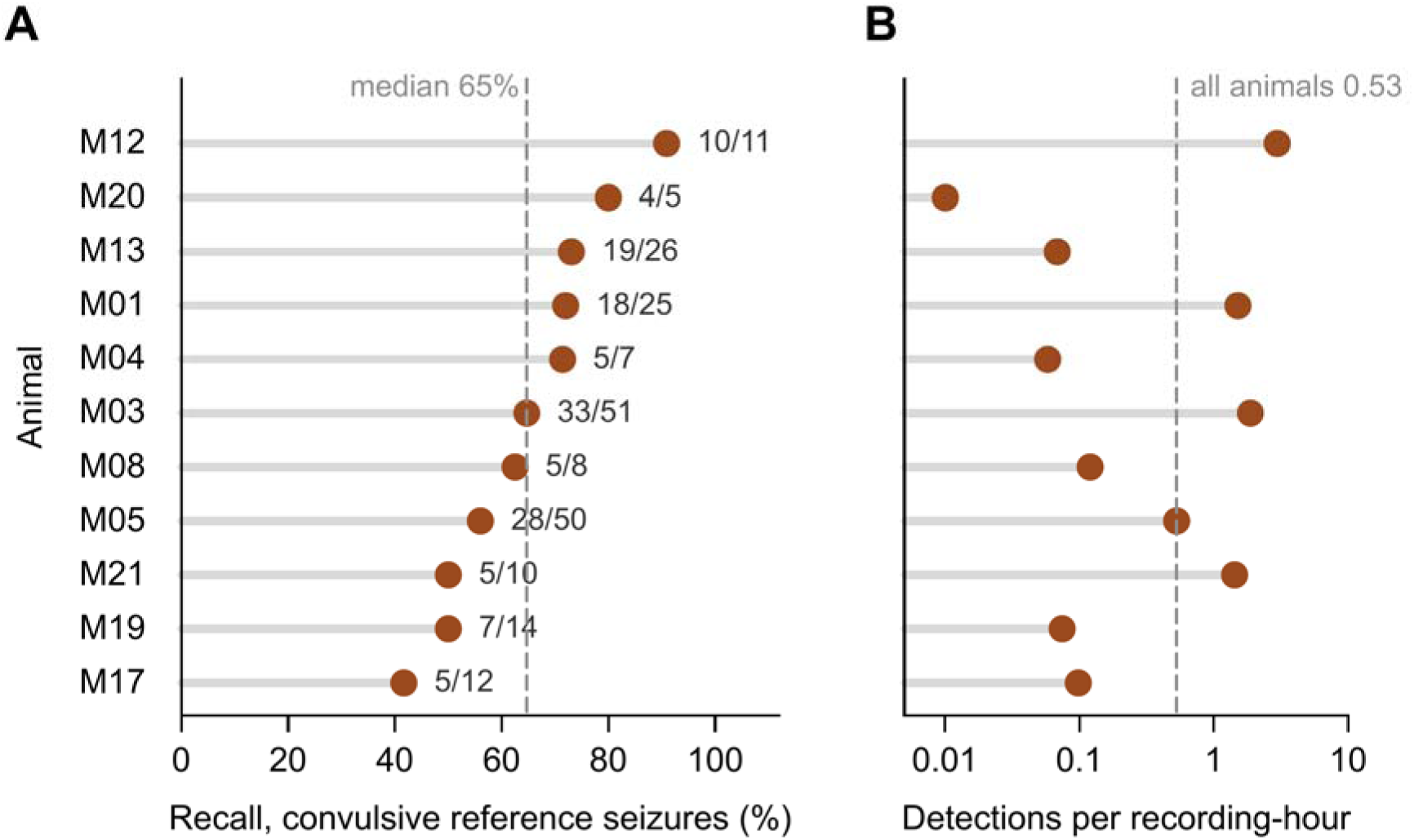
The source detector detects most convulsive seizures in a new cohort. Second cohort, Batches 1 to 3 (7,142 recording-animal units, 10,461 h), 221 video-adjudicated convulsive reference seizures. A reference seizure counts as detected when a detection overlaps it within 5 s. The source detector was trained on the first cohort only and run at a detection threshold of 0.5 and a boundary threshold of 0.1. (A) Recall for the 11 second-cohort animals with at least five reference seizures. Numbers give detected / reference seizures.

**Table 1.** The source detector on its own cohort and on the second cohort. Note that the difference in detection rate depends on cohort seizure frequency, not detector performance.

|  | **Source cohort** | **Second cohort** |
|---|---|---|
| **Training** | 867 confirmed seizures, 165 recordings | none |
| **Recall** | 93% (380/408) on three animals whose seizures were not used in training (validation F1 0.83 at threshold 0.5) | 64% (141/221) of video-adjudicated convulsive seizures |
| **Per-animal recall** | 93% (325/348) and 92% (55/60) in the two animals with seizures; the third had no seizures and no detections (21 recordings) | median 65%, range 42–91% (11 animals), none at zero |
| **Detections per recording-hour** | 3.6 (689 in 191 h), 2.5 h flagged | 0.53 (5,544 in 10,461 h), 17.6 h flagged |
| **Precision** | 96% (661 of 689 detections), on the same three animals | 72% (90 of 125 detections) in 55 held-out recordings, where 19 of 30 reference seizures were detected |

Normalization affects how the detector transfers. A control trained on the same annotations with the same code, but with each 60-s window standardized on its own, produced five times as many detections at the same operating point (29,258, 2.80 per hour, 73.9 h flagged, for 189 of 221 seizures), and at a matched number of detections it recalled 37 reference seizures where the per-recording model recalled 88 (Supplementary Table 3).

Dashed line, median across animals (65%, range 42–91%). Overall, the detector found 141 of 221 reference seizures (64%). (B) Detections per recording-hour for the same animals (log scale, about 500 h per animal). Dashed line, all recordings (5,544 detections in 10,461 h, 0.53 per hour). Data in Supplementary Table 2.

## Annotating the new cohort in the app and retraining improves detection

The second cohort was annotated in the review queue: 344 detector proposals were adjudicated, and 48 events were added manually. Together with the video-adjudicated reference, these became the second cohort’s training labels. Review in the queue took a median of 2 to 7 s per proposed event in the four review sessions we timed, 349 events in about 38 min of labeling (Methods). The detector was retrained on the source cohort’s annotations plus these, holding out each animal’s latest recordings so that every animal was represented in training (Methods).

Scored on the same recordings against the same reference, recall rose from 141 to 195 of 221 (88%), with 8,190 detections (0.78 per hour) and 29 h flagged (Table 2). Per-animal recall rose in 10 of the 11 animals and was unchanged in one (M20, 4 of 5 before and after), from a median of 65% to 92% (Wilcoxon signed-rank test, P = 0.002, Fig. 4A). On the held-out units the retrained model detected 27 of the 30 reference seizures against 19 for the source model (Fig. 4B), and 142 of its 193 detections there were seizures (74%, Fig. 4C), so that retraining found 142 seizures where the source model had found 90, at the same precision (74% against 72%). Detections also matched the reference seizures more closely, with a median intersection over union of 0.70 against 0.58 (Supplementary Table 1). Precision varied between animals: all detections were seizures in seven animals, and the three animals with most detections were at 83% (38 of 46), 78% (29 of 37) and 73% (29 of 40), with two animals at 38% (11 of 29) and 29% (2 of 7).

**Figure 4.**
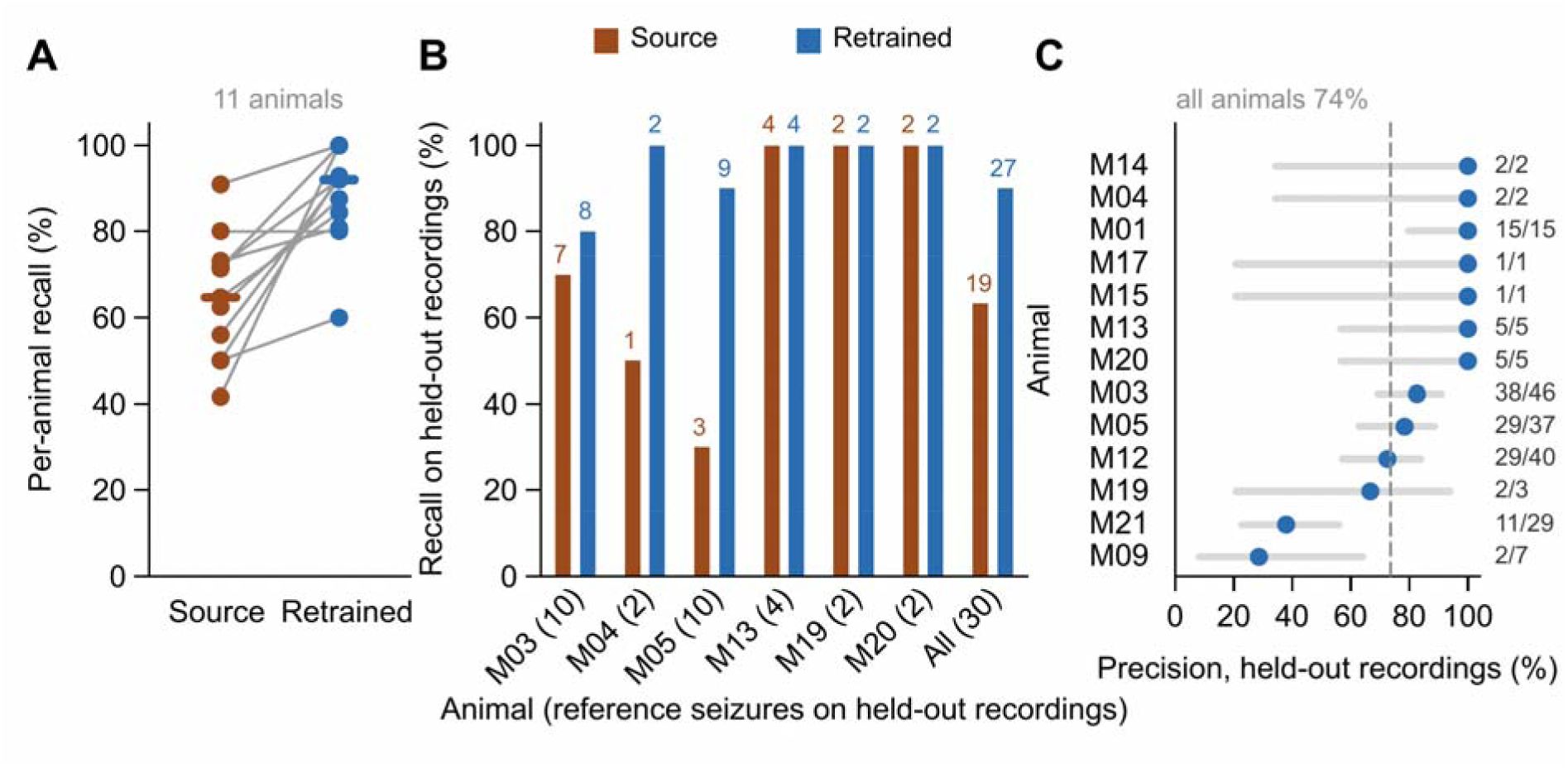
Retraining with annotations from the new cohort improves detection. Source detector, trained on the first cohort only. Retrained detector, trained on the first cohort plus the second- cohort annotations made in the app. Both at detection threshold 0.5 and boundary threshold 0.1. (A) Recall per animal for the two detectors (same 11 animals as Fig. 3, lines join the same animal, bars mark the medians, 65% and 92%). Recall rose in 10 animals and was unchanged in one (Wilcoxon signed-rank test, P = 0.002). Over all animals, 141 and 195 of 221. These recordings include those the retrained detector was trained on. (B) Recall on held-out recordings, the 55 recording-animal units that the retrained detector did not train on (it did train on earlier recordings of the same animals), for the six animals with reference seizures there and for all (number of reference seizures in brackets, numbers above bars give seizures detected): source 19 of 30, retrained 27 of 30. (C) Precision of the retrained detector on the held-out recordings, per animal: share of detections that are seizures after review of every detection (numbers, seizures / detections, grey bars, 95% Wilson intervals). Dashed line, all animals (142 of 193, 74%, against 90 of 125, 72%, for the source detector).

**Table 2.** NED-Net and SeizyML on the second cohort. Both tools trained on the first cohort only (source) and retrained with the same second-cohort annotations, scored on the same recordings against the same reference with the same matcher. SeizyML scored at its automatic stage, without the manual verification its workflow intends.

|  | **NED-Net source** | **NED-Net retrained** | **SeizyML source-trained** | **SeizyML retrained** |
|---|---|---|---|---|
| **Reference seizures detected (of 221)** | 141 (64%) | 195 (88%) | 151 (68%) | 146 (66%) |
| **Detections (per recording-hour)** | 5,544 (0.53) | 8,190 (0.78) | 17,323 (1.66) | 10,809 (1.03) |
| **Recording time flagged** | 17.6 h | 29.0 h | 220 h | 132 h |
| **Held-out reference seizures detected (of 30)** | 19 | 27 | 22 | 22 |
| **Held-out detections that were seizures** | 90/125 (72%) | 142/193 (74%) | 58/138 (42%) | 55/117 (47%) |

## Comparison with a published detector on the same recordings

We then compared NED-Net with SeizyML^5^, a published rodent seizure detector with a graphical interface and open code, used as its authors intended. Whole recordings, its own preprocessing, features and Gaussian naive Bayes classifier, and its published default thresholds. SeizyML ships no pretrained model, so we trained it twice to mirror our two models, on the 165 source-cohort recordings and their annotations, and on these plus the second-cohort recordings and annotations the retrained NED-Net model was trained on (Methods). Both were run on the same recordings and scored against the same reference with the same matcher (Fig. 5).

**Figure 5.**
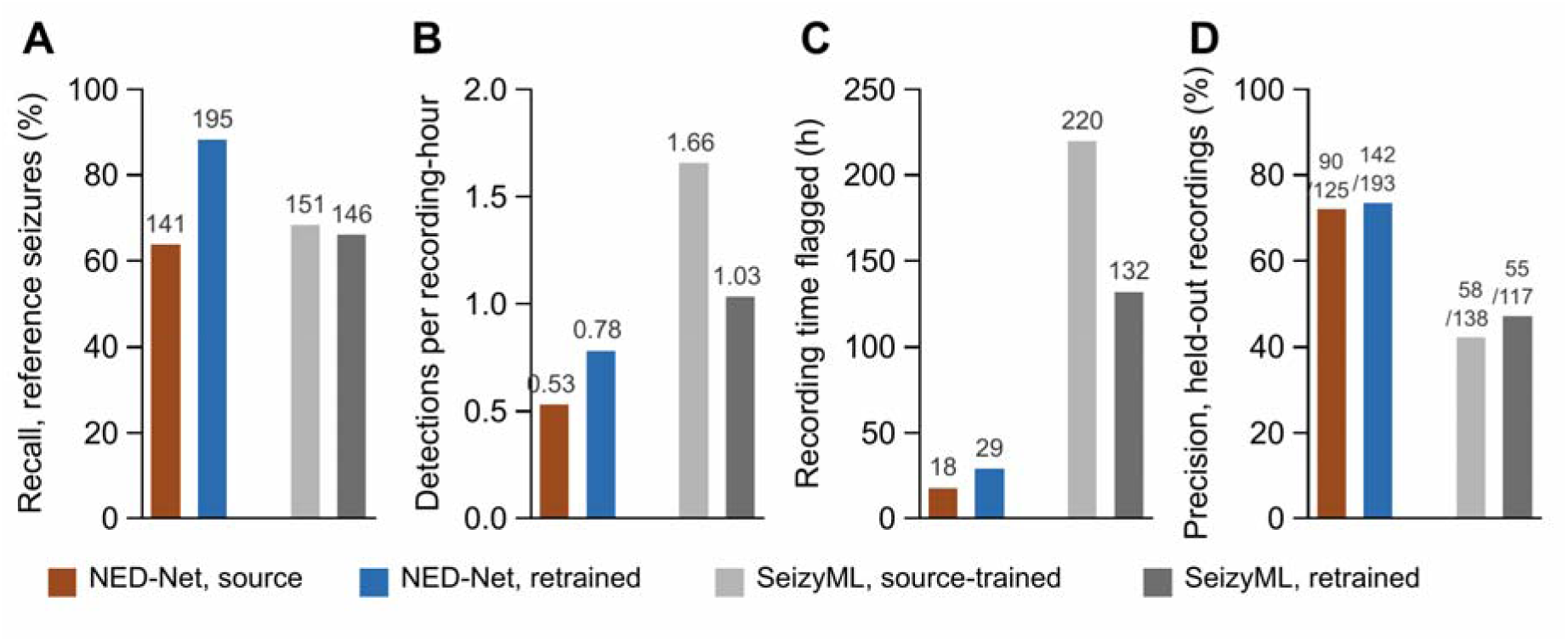
Comparison with SeizyML. NED-Net and SeizyML (Antonoudiou et al. 2025), each trained on the first cohort only ("source") and retrained with the same second-cohort annotations, run on the same recordings and scored against the same reference. SeizyML was run on whole recordings with its published defaults, and its automatic output is scored without the manual verification its workflow intends. (A) Recall of the 221 reference seizures (numbers, seizures detected): NED-Net 141 and 195, SeizyML 151 and 146. (B) Detections per recording- hour: 0.53 and 0.78 for NED-Net, 1.66 and 1.03 for SeizyML. (C) Recording time flagged as seizure: 18 and 29 h, and 220 and 132 h. (D) Precision on the held-out recordings after review of every detection (numbers, seizures / detections): 90/125 and 142/193, and 58/138 and 55/117.

Trained on the source cohort, SeizyML detected 151 of the 221 reference seizures (68%), close to NED-Net’s 141, but with 17,323 detections (1.66 per hour) and 220 h flagged as seizure, three times the detections and twelve times the flagged time of the NED-Net source model. On the held-out units, it detected 22 of the 30 reference seizures, and 58 of its 138 detections were seizures on review (42%), against 72% for the NED-Net source model. Retrained with the same second-cohort annotations, SeizyML detected 146 reference seizures (66%) with 10,809 detections (1.03 per hour) and 132 h flagged, again 22 of 30 held out, and 55 of its 117 held-out detections were seizures (47%), whereas the retrained NED-Net model reached 195 of 221, 27 of 30 and 74%. SeizyML’s events are built on a 5-s grid and were longer than NED-Net’s (median 35 s against 8 s). Two differences in how the tools were used affect this comparison.

SeizyML is semi-automated, and its intended output is produced only after each event has been manually verified. The figures above therefore describe only its automatic stage. In addition, its training labels treated every 5-s segment outside a confirmed seizure as background, including unreviewed activity, whereas NED-Net’s training draws background windows only from stretches untouched by any annotation (Methods).

## Example detections on held-out recordings

Figure 6 shows three reference seizures on recordings neither model was trained on. In the first, both models detect the seizure within seconds of its onset. In the other two, the source model’s probability crosses the detection threshold only briefly or not at all, and the retrained model detects the seizure. The examples were chosen by rule, one per animal (Methods).

**Figure 6.**
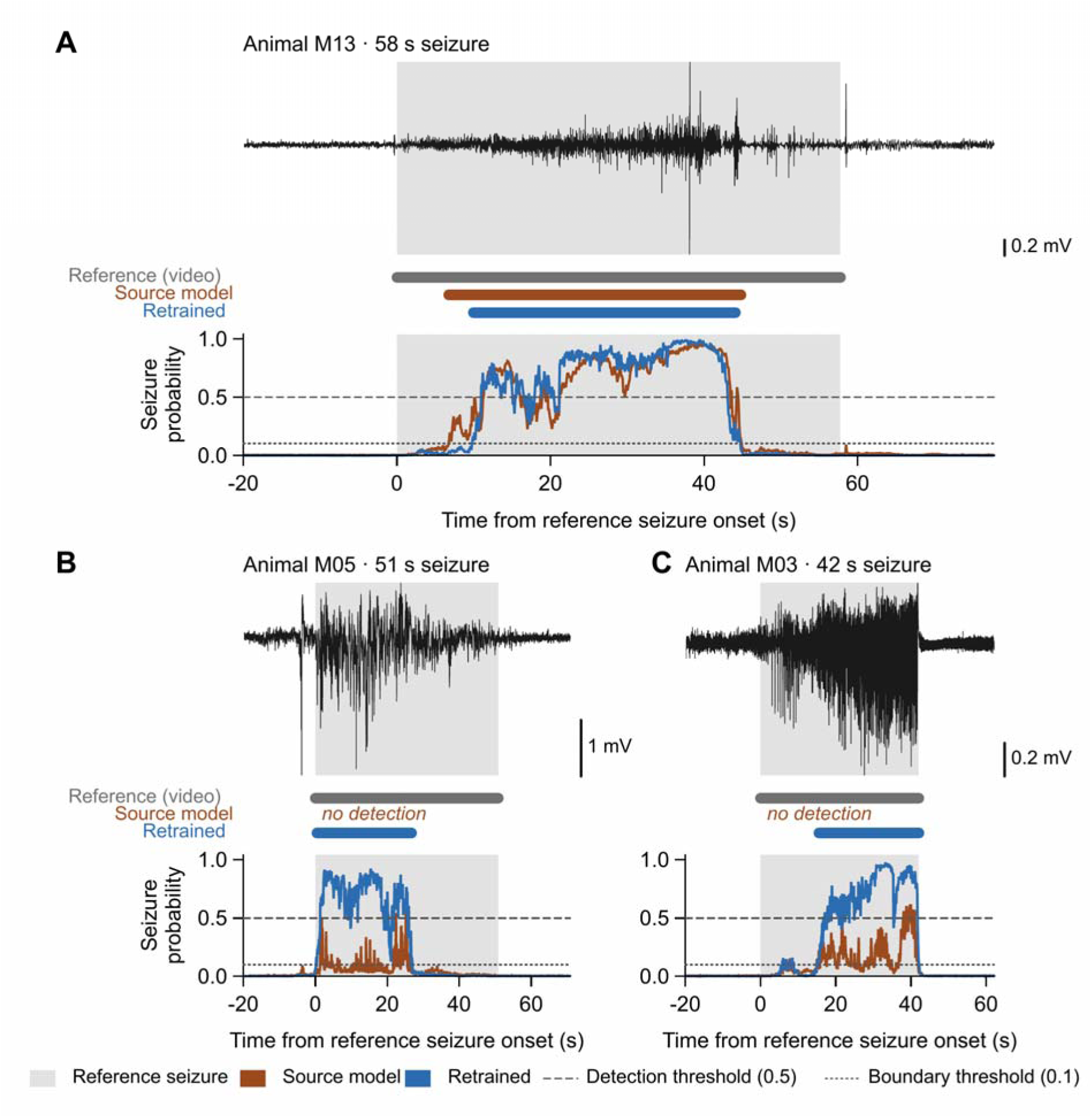
Example detections on held-out recordings. Three reference convulsive seizures (grey shading) from three animals, on recordings that neither detector was trained on. In each panel, the top trace is the EEG, the middle row shows the reference seizure and each detector’s detections, and the bottom row shows each detector’s seizure probability with the detection threshold (0.5, dashed) and the boundary threshold (0.1, dotted). (A) A 58 s seizure detected by both detectors (source from 7 to 45 s after the reference onset, retrained from 10 to 44 s). (B) A 51 s seizure missed by the source detector, whose probability passes 0.5 only briefly, under the 5 s minimum duration, and detected by the retrained detector from 0.5 to 27 s. (C) A 42 s seizure missed by the source detector and detected by the retrained detector from 16 to 42 s.

Examples were chosen by rule, one per animal: the longest held-out seizure detected by both detectors, and the two longest held-out seizures, in two further animals, missed by the source detector and detected by the retrained one.

## A convulsive classifier removes non-seizures and enriches convulsive seizures

Detection in NED-Net is a cascade. The detector (Stage 1) proposes events, and a second network (Stage 2) assigns each event a convulsive probability without adding or removing events, so that a user can count convulsive seizures separately or review them first. We trained the classifier on the confirmed seizures of both cohorts, split by convulsive flag, and applied it to the retrained detector’s output (Fig. 7). Its operating point, 0.8, was the threshold that maximized event-level F1 on its validation data. Over all second-cohort recordings, the classifier traded recall of the convulsive reference for precision against it: 195 of 221 (88%) at 9% with no filter, 133 (60%) at 24% at threshold 0.65, 70 (32%) at 39% at 0.8, and 18 (8%) at 82% at 0.95 (Fig. 7A). On the held-out units, where every detection had been reviewed, 142 of the retrained detector’s 193 detections were seizures (74%) and 37 were convulsive (19%). At threshold 0.65, 53 of the 58 detections that passed were seizures (91%) and 27 were convulsive (47%), with 22 of the 30 reference seizures kept. At 0.8, all 26 detections that passed were seizures, 19 of them convulsive (73%), with 17 of 30 reference seizures kept (Fig. 7B). What the classifier does on this cohort is remove the non-seizures from the detector’s output and raise the convulsive share of what remains from 19% to 73%. It does not make every detection convulsive, and the choice of threshold sets how many convulsive seizures are kept.

**Figure 7.**
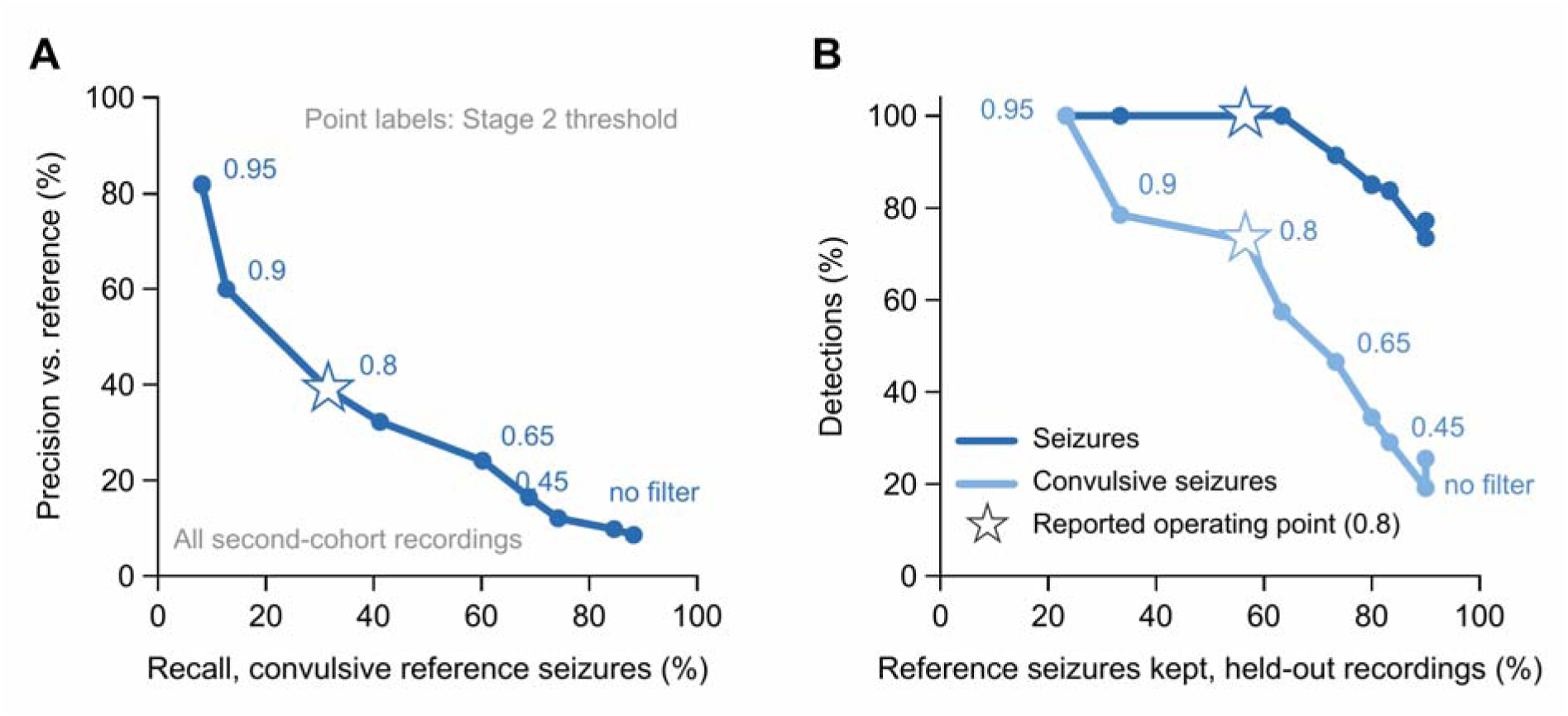
The convulsive classifier removes non-seizures and enriches convulsive seizures. Stage 2 classifier applied to the retrained detector’s detections. Point labels give the Stage 2 threshold ("no filter", detector alone), and the star marks the reported operating point (0.8, the threshold with the highest F1 on the classifier’s validation data). (A) All second-cohort recordings: precision against the reference (among detections overlapping an event the video reviewer adjudicated, the share that are reference seizures) against recall of the 221 reference seizures. No filter, 195 of 221 (88%) at 9%. At 0.65, 133 (60%) at 24%. At 0.8, 70 (32%) at 39%. At 0.95, 18 (8%) at 82%. (B) Held-out recordings: share of detections that are seizures (dark) and that are convulsive seizures (light, overlapping a reference seizure or marked convulsive by the reviewer), against the share of the 30 reference seizures kept. No filter, 142 of 193 seizures (74%), 37 convulsive (19%), 27 of 30 kept. At 0.65, 53 of 58 (91%), 27 (47%), 22 kept. At 0.8, 26 of 26 (100%), 19 (73%), 17 kept.

## Thresholds do not transfer between models

The operating point of a retrained detector does not carry over from the model it replaces. When an earlier retrained model was run at the source model’s thresholds, it labeled 72% of all recorded time as ictal, with a median event of 1,713 s, and reached an apparent convulsive recall of 69%, close to what that coverage alone would produce. At its own operating point, the same model labeled 0.01% of recorded time. Output-score distributions change after retraining, and a published threshold applied to retrained weights can produce a figure that resembles high sensitivity. Each model reported here was therefore assigned the threshold with the highest event-level F1 on its own validation data, and the released software stores an operating point with each model and applies it when the model is selected, so that thresholds are not carried over by default to weights they were not validated on.

## Discussion

We describe NED-Net, open-source software in which annotation, review, training, and detection of rodent EEG events share one data structure, and we evaluated it on two cohorts of kainate-treated mice. A detector built in the software on the first cohort detected 64% of the second cohort’s video-adjudicated convulsive seizures as delivered, left no animal without a detected seizure, and 72% of its detections on held-out recordings were seizures. Annotating the second cohort in the review queue, at a few seconds per event, and retraining raised recall to 88% and held-out recall from 19 to 27 of 30 at the same precision. A published detector trained on the same labels reached similar recall at the start with three times the detections and did not gain from retraining. A convulsive classifier removed every non-seizure from the retrained detector’s held-out output at its validation-chosen threshold, with 73% of what remained convulsive.

The observation that a detector improves when trained on annotations from the experiment at hand is not itself a finding, but the premise on which the software is built. The measurements reported here quantify that improvement: 54 additional detections among 221 reference seizures and 8 among 30 held-out seizures, across 392 events reviewed or added to the queue. This premise depends on annotation being inexpensive and detection running early enough for annotations to inform the analysis. The software is designed to meet both requirements. A review queue lets users confirm or reject proposed events in seconds; detection runs on the acquisition folder as files are written; and per-animal reporting is available from the first file.

Variation between cohorts and animals has been reported previously^2,3,4^, and cross-subject variability in seizure morphology has been identified as the main obstacle to generalization, even for spike-wave discharges^6^. We observed this variability as well, as per-animal recall ranged from 42% to 91% before retraining and from 60% to 100% afterward. Accordingly, the software attributes every event to an animal and reports results per animal, and we recommend reviewing each animal before relying on any detector’s output.

The way the signal was scaled shaped how the source detector transferred to a new, unseen cohort. A control trained on the same labels with the same code, with each 60-s window standardized independently, ranked events less well: at a matched number of detections it recalled 37 of the 221 reference seizures where the detector using a center and scale computed once for each recording and channel recalled 88, and at the same operating point it needed five times as many detections (Supplementary Table 3). One plausible explanation is that amplitude varies more across implants than within a recording, because of animal-to-animal variability in electrode position. Standardizing each window independently discards its amplitude relative to the rest of the recording, which may help distinguish seizures from background on an unfamiliar implant. Other detectors that do not retrain also use forms of per-implant scaling: the feature- based detector of Kamintsky et al.^1^ normalizes each feature against a manually selected seizure-free baseline segment from each recording; SeizyML normalizes features per recording by design^5^; and early patient-specific clinical detectors were trained on each patient’s own seizures because within-subject consistency exceeds between-subject consistency^7^. These results suggest that how a detector handles signal scale across implants is a first-order design choice when transferring between cohorts.

The comparison with SeizyML should be understood in light of its design. SeizyML supports a workflow where every detected event is manually verified, and its authors describe it as semi- automated. In these recordings, it detected a similar proportion of the convulsive reference seizures as the NED-Net source model, but flagged three times as many events and twelve times as much recording time. Fewer than half of its detections on the held-out data were seizures, so users would spend correspondingly more time verifying them. Its failure to benefit from annotations in the second cohort is consistent with a low-capacity classifier using fixed features having little to learn from additional labels of the same kind. However, some of the difference may reflect the training labels, as SeizyML treated unreviewed activity as background. NED-Net adds a detector that learns from user annotations, identifies shorter events that require less recording time to review, and includes a convulsive classification stage that the comparator lacks.

The numbers after retraining also show what the loop did not accomplish. Recall of 88% and precision of 74% are adequate for a detector whose output is reviewed, as the software is intended to be used. They are not adequate for unattended counting of convulsive seizures, however, and performance still varied between animals: one animal remained at 60% recall, and two animals had precision below 40%. We did not systematically measure signal quality, but the cohort required excluding one animal because of electrode failure and two channels because of persistent artifact. In our experience, recording quality and electrode placement are the first things to address, before the model. The Viewer tab is designed for this purpose: it displays a full-length recording alongside its detections, activity channel, and synchronized video, allowing users to assess each channel’s quality and the plausibility of detections across hours rather than from isolated events. Beyond that, the results suggest three steps. First, add annotations for the animals identified as weak in the per-animal report. Second, run the classifier at a threshold suited to the task: at 0.65, nine of ten passing detections are seizures and 22 of 30 convulsive reference seizures are retained, whereas the validation-selected threshold of 0.8 passes only seizures but retains 17. Third, fine-tune the detector for each animal, as with patient-specific clinical detectors. The sidecar structure supports this because every event is already attributed to an animal.

The software presents model output as a review queue. A channel on which the detector is silent can be identified automatically and routed to review or to the rule-based detectors, but a channel on which it fires steadily but wrongly produces a plausible event count that is only exposed by a reviewer. Confidence-scored output routed to a human has been used before in rodent EEG, for spike-wave discharges^8^ and in SeizyML^5^. NED-Net differs in that validated events are not the endpoint, but the training set for the next model, and in attributing every event to an animal so that review can target the implants where the model is weak.

Live detection changes when the loop can run. Real-time seizure detection in rodents has so far been developed for closed-loop intervention, in which detection triggers a stimulus within seconds^9,10^. Our goal is different, to make the detector’s behavior on each implant visible while the experiment is still running. The measurements reported here were collected after acquisition had ended, but in the intended use, per-animal output is available within the first few days of a recording protocol, allowing annotation effort to focus on animals that need it before most of the protocol has been recorded.

Neither machine learning nor the seizure phenotype is required for the workflow, and the software is intended as a complete analysis path rather than only a model. The rule-based detectors, the review queue, and the per-animal aggregation need no trained model or GPU, in the tradition of the signal-processing detectors that preceded machine learning in this field^11^, and in our own use they produced the first annotation set from which we trained the first model. From there, a laboratory can train on a single GPU, run detection unattended on a cluster or live on the acquisition folder, convert LabChart recordings to EDF, and keep every annotation with its provenance in a sidecar next to the recording it belongs to, so that no annotation is lost when a model is replaced. The Results tab reports counts, durations, and seizure burden per animal and per group. The same machinery also serves an interictal spike detector, and the models can be trained on any event type a reviewer marks as confirmed or rejected. Among published rodent detectors with a graphical interface^1,5^, NED-Net is, to our knowledge, the only one that closes the loop from review back to training, learns from the user’s own annotations rather than fixed engineered features, and runs during acquisition.

Finally, how validation events are selected affected apparent performance more than any other choice we examined. In this work, the same detector output 0.12% precision when all detections were pooled without a threshold, 100% when files were chosen for high confidence, 34% in a random sample of events, and 92% when the detector fired in a review of whole recordings.

That evaluation methodology can move apparent performance by this much has been shown for clinical detectors^12^, and the SzCORE framework was proposed because heterogeneous validation made published algorithms incomparable^13^, with re-implementation under standardized conditions revealing substantial discrepancies from reported performance^14^. To our knowledge, no equivalent standard exists for rodent EEG, where seizure rates, electrode configurations, and reference annotations vary widely. We therefore chose every operating point by one rule applied to validation data, measured precision by reviewing every detection on held- out recordings rather than a sample, and ran the comparison tool under a protocol written before its output was scored, and we argue that detector validation in this field should state its sampling scheme with the same care as its model architecture.

## Methods Software

NED-Net (Neural Event Detection Network) is written in Python (≥3.9) using the Dash and Plotly libraries. It runs as a local web server and can be opened in a browser or in a desktop window using pywebview. Installers for standalone versions are also available. EDF files are read with pyEDFlib, and LabChart (AD Instruments). adicht recordings can be converted to EDF with a built-in tool. Detection, training and inference are implemented in a single package that is shared between the graphical interface and the command-line entry points, which also run unattended under SLURM on a computing cluster. The source code is released under the GNU General Public License version 3 at https://github.com/EpilepsyCenter/NED-Net.

The central data structure is a sidecar annotation file (

<RECORDING>_ned_annotations.json) stored next to each EDF recording. Each event records its channel, onset and offset, a label (confirmed, rejected or pending), the method and the tab or mode that proposed it (manual annotation, rule-based detection, or single-file, batch or live analysis), and a features dictionary containing the detector confidence and, when the convulsive classifier has run, the convulsive label and its probability. The model outputs pending events, which populate the review queue, and reviewed events are written back as confirmed or rejected, which become the training labels for the next model. A second sidecar assigns each EEG channel to an animal, cohort and experimental group. In the recordings analyzed here, each file contains eight animals, one per channel.

The interface is organized in functional tabs (Fig. 2, Supplementary Fig. 1). The Load and Viewer tabs open one or more EDF files, pair EEG channels with activity channels recorded at a different sampling rate (if applicable), apply or edit the channel-to-animal map, and display filtered multi-channel traces with existing detections, an activity overlay and synchronized video. The Detection tab provides four rule-based seizure detectors that require no trained model (spike-train, spectral band-power, autocorrelation, and an ensemble of these by voting), each with configurable thresholds, minimum duration and merge gap, followed by boundary refinement, quality scoring and optional post-detection filters. The same annotation and training workflow also supports an interictal spike detector. The Training tab builds datasets from the sidecars, trains the models described below with the signal scaling of choice (per recording by default), and hosts the review mode, a queue of pending events ordered by confidence or duration in which the reviewer inspects the trace around each event with adjustable context, an amplitude overview of the whole recording and keyboard navigation between detections, may adjust its boundaries, and confirms or rejects it. Decisions are written to the sidecar immediately and manual events can be added by dragging on the trace in Browse mode. The Analysis tab runs the trained cascade on a single file, on a folder, or live during acquisition. Selecting a bundled model fills in the operating point validated for it, and other models default to the optimum found at training. The Results tab aggregates every run in a per-project SQLite database and reports counts, durations and seizure burden per animal and per group. Any run’s detections can be reviewed from the Results tab in the same confirm/reject queue as training proposals. Rejected detections are removed from the counts and burden figures, and confirmed and rejected labels are written to the sidecars, where they become training material for the next model.

In live mode the software monitors an acquisition folder. When a new file appears, it waits for a configurable delay (default 30 s) to allow the recorder to finish writing the file, then runs the full cascade, and writes the resulting events to the file sidecars. The channel-to-animal map for live mode is supplied as a session template keyed by channel rather than by filename. Since the channel-to-animal mapping is fixed for a recording session, we attribute every event to an animal from the first file onward. Files recorded before monitoring started can be processed as a backlog.

## Models

The detector (Stage 1) is a one-dimensional U-Net^15^ that receives 60-s single-channel windows sampled at 250 Hz and returns a per-sample seizure probability. Before windowing, each channel of a recording is normalized with a center and scale computed once over the whole recording, the median and 1.4826 times the median absolute deviation, and clipped at ±30 (per- recording normalization). The statistics are stored with each model and applied automatically at detection, and standardization of each window to zero mean and unit variance is retained as an option. The encoder consists of four levels of two convolutional blocks each (kernel size 7, batch normalization, ReLU) with 2× max-pooling and a doubling of channels at each level from 32 base filters, and the decoder mirrors it with transposed convolutions and skip connections (dropout 0.2, 23.4 million parameters). The network is trained with a combined Dice and binary cross-entropy loss, using AdamW (learning rate 10⁻³, weight decay 10⁻, batch size 8, a weight of 5 on the positive class in the cross-entropy term), a learning-rate schedule that reduces the rate when the validation loss plateaus, and early stopping (patience 10), in full fp32 precision.

Events are extracted from the probability trace by hysteresis thresholding, with a detection threshold that an event must reach and a lower boundary threshold that determines its extent, followed by a minimum duration (5 s) and a merge gap. The convulsive classifier (Stage 2) uses the same convolutional encoder followed by global average pooling, dropout (0.3) and a linear output layer, and returns one convulsive probability per 60-s crop centered on the event, with each crop standardized on its own. It is trained with bf16 mixed precision (learning rate 3 × 10⁻, batch size 16, 30 epochs, positive weight equal to the non-convulsive/convulsive ratio). Stage 2 writes a convulsive flag and probability to the event but never adds, removes or filters events, so the set of detected events depends on Stage 1 alone.

## Animals and recordings

All experimental protocols involving animals were reviewed and approved by the Malmö/Lund Ethics Committee on Animal Testing at the Lund District Court, Sweden (Permit No. 2721- 2025), under the authority of the Swedish Board for Agriculture (Jordbruksverket), in accordance with the Swedish Animal Welfare Act and EU Directive 2010/63/EU.

### Source cohort

Adult male C57BL/6 mice (8–12 weeks, Janvier Labs) were rendered chronically epileptic by inducing status epilepticus (SE) with repeated low-dose subcutaneous injections of kainic acid (5 mg/kg)^16^. Six weeks after SE, mice were anesthetized with isoflurane (3–4% for induction and 1–2% for maintenance) and positioned in a stereotaxic frame. A two- lead biopotential telemeter (Kaha Sciences, AD Instruments) was implanted in a subcutaneous pocket on the right side of the back. One lead was connected to a stainless-steel recording electrode implanted in the right hippocampus at AP −2.2 mm, ML −1.7 mm and DV −2.0 mm.

The second lead was connected to a reference electrode positioned over the contralateral dura using a stainless-steel screw. The electrode assembly was secured with dental cement, and the incision was sutured. Animals were allowed at least two days of postoperative recovery before the start of EEG recording. Each animal’s cage was positioned over a dedicated tBase (Kaha Sciences), which wirelessly powered the implanted telemeter and received its signals. EEG signals were acquired through PowerLab (AD Instruments), sampled at 2000 Hz, and recorded using LabChart Pro (AD Instruments). Recordings consisted of three batches of eight mice for six weeks each. Of these, we annotated 165 recordings in the interface: we reviewed rule-based detections and added manual events, yielding 1,136 human-reviewed events, of which 867 confirmed seizures (maximum duration 84 s) formed the training positives for the source model.

### Second cohort

Twenty-four additional male C57BL/6 mice were treated as above, implanted and recorded in three batches of eight on the same acquisition system. A fourth batch of the same experiment was not analyzed. This gave 1,020 recordings of up to 92 min. None of these recordings were included in the source model’s training set. One Batch 1 animal was excluded for electrode failure and two Batch 2 channels for persistent recording artifact, leaving 21 animals, 7,142 recording-animal units and 10,461 recording-hours on which all figures are reported. Animals are identified by neutral codes (M01 to M21) in the text, figures and deposited data.

### Euthanasia

At the end of the experimental period, all animals were given an intraperitoneal overdose of sodium pentobarbital and transcardially perfused first with 50 ml 0.9% saline and then with 50 ml of ice-cold 4% paraformaldehyde (PFA). Brains were extracted, post-fixed overnight in PFA, cryo-protected in 30% sucrose solution in 0.1% phosphate buffer, and finally sliced into 30 µm slices with a freezing microtome. Sections were stored in a glycerol and ethylene glycol solution (20% and 30% respectively, in 0.1% phosphate buffer) at −20 °C until further processing.

## Reference annotations

Two annotation sets served different purposes. The first is an external, video-adjudicated convulsive reference. A classical candidate generator implemented in SETA (Seizure Event Tagging and Annotation), which detects 15–80 Hz envelope peaks above median + 4 × MAD and groups them into episodes of at least 15 s at a density of at least 2.5 spikes/s, proposed candidate events across the three second-cohort batches, and each candidate was adjudicated on synchronized video by a reviewer without access to NED-Net output. On the retained channels this yielded 221 convulsive or behavioral seizures, 68 in Batches 1 and 2 and 153 in Batch 3. Because this reference contains no non-convulsive events, which video cannot reliably confirm, it measures convulsive recall but cannot credit a non-convulsive detection. Precision measured against it ("precision vs. reference") counts, among detections that overlap an adjudicated candidate, the share that overlap a confirmed convulsive seizure. The second set comprises the annotations made in the interface during review of the second cohort, 344 adjudicated detector proposals and 48 manual additions, which are not independent of the detector and were used as training data but not as a recall reference.

## Training sets and retrained models

Positives were confirmed events from the sidecars, capped at 100 s. Hard negatives were windows centered on rejected events, excluding rows the video reviewer had rejected, because a rejection in that reference means "not a convulsive seizure" rather than "not a seizure" and these rows sometimes contained non-convulsive activity. They were retained in the training tree so that background sampling would avoid them. Background negatives were random windows drawn from annotated recordings and never overlapping an annotated region of either label, at six negatives per positive. The source detector was trained on the source-cohort annotations alone (2,884 training and 1,451 validation windows, validation on animals held out from training). The retrained detector was trained on the source-cohort annotations, the external reference’s confirmed seizures for the second cohort, and the annotations made in the interface during review of the second cohort, with the same recipe (8,245 training and 1,177 validation windows). Its split held out each animal’s latest recordings until 20% of that animal’s positive windows were withheld, so that every animal was represented on both sides of the split, which corresponds to the intended workflow, in which the early recordings are annotated and the model handles the remainder. The held-out side of this split, 55 recording-animal units from 11 animals holding 30 reference seizures, is the held-out set used for precision and held-out recall throughout. Individual events were never divided between training and validation, because events within one recording share electrode state, ambient noise and frequently one seizure cluster. The convulsive classifier was trained on the confirmed seizures of both cohorts, split by convulsive flag, with rejected events as additional non-convulsive negatives, and with each 60-s crop standardized on its own, because per-recording normalization did not improve the classifier on validation data.

Checkpoints were selected on event-level F1 at the optimal threshold on the validation split, and each model’s operating point is the threshold with the highest event-level F1 on that split: 0.5 for both detectors, with a boundary threshold of 0.1 and a minimum duration of 5 s, and 0.8 for the classifier (validation F1 0.84, precision 0.87, recall 0.81). Training ran on a single NVIDIA A100 GPU at LUNARC (Lund University). Each detection pass over the 1,020 recordings took about 3 h on one 48-core CPU node.

## Comparison tool

SeizyML^5^ version 2.0.0 (commit f0d4b26) was run as its authors intend: whole recordings cut into 5-s segments at 100 Hz, its own preprocessing, features, feature selection, Gaussian naive Bayes classifier and post-processing, all at the published defaults (dual threshold 0.5 and 0.2, rolling window 6, dilation 2, erosion 1). The only setting changed was the channel configuration, one EEG channel per animal instead of two. SeizyML ships no pretrained model. Its source- trained model was trained on the 165 source-cohort recordings and annotations of the NED-Net source model (458 recording-animal units), and its retrained model on these plus the second- cohort recordings that the retrained NED-Net model was trained on, with its held-out units excluded (3,113 recording-animal units, 3.36 million segments, 4,996 seizure segments). A segment was labeled seizure when at least half of it lay inside a confirmed seizure, and every other segment, including unreviewed activity, was labeled background, which differs from NED- Net’s training, in which background windows never overlap an annotated region. SeizyML’s events were kept as produced, including events shorter than 5 s. The evaluation (recordings, reference, matcher, operating points) was written down before any SeizyML output was scored against the second cohort, and SeizyML was not tuned on the second cohort.

## Evaluation

We considered a detection a match to a reference seizure if both were in the same file and channel and their intervals overlapped, allowing a 5 s tolerance at each end. Recall was the fraction of reference seizures matched by at least one detection. Detections per recording-hour and recording time flagged as seizure (the summed duration of all detections) were computed over the 7,142 retained recording-animal units. In a sensitivity analysis, we recomputed recall without tolerance, using an intersection-over-union of at least 0.2, requiring the detection to cover at least 50% of the seizure, and using greedy one-to-one assignment (Supplementary Table 1). We computed per-animal recall for the 11 animals with at least five reference seizures. The example seizures in Fig. 6 were chosen by rule, one per animal: the longest held-out reference seizure detected by both models, and the two longest held-out reference seizures, in two further animals, missed by the source model and detected by the retrained model.

Precision was measured on the 55 held-out recording-animal units by review of every detection, none skipped. Four reviews contributed labels: every detection of an earlier retrained model on these units (141 events), every detection of the source-trained SeizyML model (138 events), and top-up reviews of every detection of the source and retrained NED-Net models that no earlier review had covered (53 and 27 events). All reviews were made by the same reviewer in the interface on the full recordings, each event confirmed when a seizure lay within it, with a convulsive flag set when the seizure was convulsive. Labels were pooled per event, and a detection counts as a seizure when the pooled labels confirm it or it overlaps a reference convulsive seizure. A detection confirmed in one review and rejected in another counts as not a seizure. The reviewer was not blind to which tool produced an event, since SeizyML’s events sit on a 5-s grid. The pooled labels hold 170 known seizures on these units. Review speed was measured from the timestamps written to the sidecar at each decision, within each review session, excluding the time before the session’s first decision.

Precision of the source detector on its own cohort was measured on the three source-cohort animals whose seizures were not used in its training, the animals of its validation split (129 recording-animal units, 191 h, 408 confirmed seizures). Of its 689 detections there, 386 lay on seizures confirmed in earlier review. A random sample of 100 of the remaining 302 detections was reviewed in the interface, 91 of which were seizures, and precision was estimated as the 386 confirmed plus 91% of the 302, that is 661 of 689 (96%), with a 95% interval of 93–97% from the sampled share. This design differs from a check on selected recordings: the animals’ seizures were not in the model’s training data, and part of the estimate comes from a sample. Nearly all of these detections come from two of the three animals, since the third produced almost none.

## Statistics and reproducibility

Counts and percentages are reported with their denominators. 95% confidence intervals in the figures are Wilson intervals on counts. The change in per-animal recall between the source and the retrained detector was tested with a two-sided Wilcoxon signed-rank test over the 11 animals with at least five reference seizures. No statistical method was used to predetermine sample size. The cohorts were those available in the laboratory, and the exclusions are stated above. The video reviewer who constructed the reference was blinded to detector output. No randomization was used, as the study compares analysis methods on the same recordings.

Each model was trained once (one seed). All numbers in the paper are recomputed from the deposited detections and labels by the scripts in the data deposit.

## Use of large language models

The authors used large language models (Claude, Anthropic) to assist with code development and analysis scripting under their direction. The authors verified all analyses and results and take full responsibility for the content.

## Data availability

The dataset needed to reproduce the results is deposited at Zenodo, DOI 10.5281/zenodo.23315437 (version 2, about 3 GB, CC BY 4.0). It contains the trained detectors and classifier, the EEG of the 55 held-out recording-animal units as single-animal EDF files with scrubbed headers, all detections of both NED-Net models and both SeizyML models on the second cohort, the video-adjudicated reference, the pooled review labels, the training windows and splits, the SeizyML models and settings, and one table per figure panel holding the plotted values. Scripts in the deposit recompute every reported number from the stored detections and labels and re-run detection on the held-out recordings. All animals and recordings in the deposit carry neutral identifiers. The full recordings (approximately 0.5 TB) are available from the corresponding author on reasonable request. They are not needed to reproduce the reported results.

## Code availability

NED-Net is available at https://github.com/EpilepsyCenter/NED-Net under the GPL-3.0 license. The analyses reported here were run with release 2.1.0, archived at Zenodo, DOI 10.5281/zenodo.24365220, which includes per-recording normalization and the deposited models. SETA, the candidate-detection and annotation tool used to construct the video- adjudicated reference, is available at https://github.com/yaqubmir/SETA, and archived at Zenodo, DOI 10.5281/zenodo.23373095. SeizyML is available at https://github.com/neurosimata/seizy_ml.

## Acknowledgements

This work was supported by the Swedish Research Council (grant number 2022-01203 to ML), the Olle Engkvist Byggmästare Foundation (grant number 233-0293 to ML), the Segerfalk Foundation (to MYM) and the Åke Wiberg Foundation (to MYM). The computations and some data analysis were enabled by resources provided by LUNARC, The Centre for Scientific and Technical Computing at Lund University (project lu2026-2-60).

## Author contributions

ML designed and supervised the study. ML wrote the software. MYM performed SE induction, electrode implantation and EEG recordings. MYM performed the video adjudication of the reference set and the annotation review of the second cohort. ML reviewed the detections on the held-out recordings. ML and MYM analyzed the data. ML and MYM wrote the manuscript. Both authors approved the final version.

## Competing interests

The authors declare no competing interests.

## Notes

https://doi.org/10.5281/zenodo.23315437

https://github.com/EpilepsyCenter/NED-Net

### Competing Interest Statement

The authors have declared no competing interest.

### Summary of Updates

Included a comparison with another published seizure detector.

