## Supplementary Data for "NED-Net: open-source software for building and refining seizure detectors during rodent EEG experiments"

Terms used throughout. Second cohort: Batches 1 to 3, 21 animals (M01 to M21), 7,142 retained recording-animal units, 10,461 h. Reference: 221 video-adjudicated convulsive seizures. A reference seizure counts as detected when a detection overlaps it within 5 s. Source detector: trained on the first cohort only. Retrained detector: trained on the first cohort plus the second-cohort annotations made in the app. Both used per-recording normalization at a detection threshold of 0.5 and a boundary threshold of 0.1. Held-out recordings: the 55 recording-animal units that the retrained detector did not train on, holding 30 reference seizures.

Supplementary Table 1. Recall of the source and retrained detectors under five event-matching rules (reference seizures detected, of 221). The primary rule, any overlap with 5 s tolerance, is used for every number in the main text. The 5 s tolerance does not add additional seizures, one-to-one matching costs one seizure per detector, and the gain from retraining holds under every rule (54 to 64 more seizures). The last two rows reflect that detections are shorter than the reference seizures. The median intersection over union of matched seizures is 0.58 for the source detector and 0.70 for the retrained one.

| **Matching rule** | **Source detector** | **Retrained detector** |
|---|---|---|
| Any overlap, 5 s tolerance (primary) | 141 (64%) | 195 (88%) |
| Any overlap, no tolerance | 141 (64%) | 195 (88%) |
| Intersection over union ≥ 0.2 | 121 (55%) | 185 (84%) |
| Detection covers ≥ 50% of the seizure | 80 (36%) | 134 (61%) |
| Greedy one-to-one assignment | 140 (63%) | 194 (88%) |

Supplementary Table 2. Per-animal recall of the source and retrained detectors (animals with at least five convulsive reference seizures, sorted by source recall), with the source detector's detections, recording time and detections per recording-hour. Data of Figs. 3 and 4A.

| **Animal** | **Reference seizures** | **Source detected** | **Source recall** | **Retrained detected** | **Retrained recall** | Source detections | Recording-hours | Source detections per hour |
|---|---|---|---|---|---|---|---|---|
| M17 | 12 | 5 | 42% | 12 | 100% | 49 | 499.2 | 0.10 |
| M19 | 14 | 7 | 50% | 13 | 93% | 37 | 497.1 | 0.07 |
| M21 | 10 | 5 | 50% | 6 | 60% | 709 | 497.1 | 1.43 |
| M05 | 50 | 28 | 56% | 46 | 92% | 264 | 499.2 | 0.53 |
| M08 | 8 | 5 | 62% | 7 | 88% | 60 | 499.2 | 0.12 |
| M03 | 51 | 33 | 65% | 43 | 84% | 935 | 499.2 | 1.87 |
| M04 | 7 | 5 | 71% | 7 | 100% | 29 | 499.2 | 0.06 |
| M01 | 25 | 18 | 72% | 23 | 92% | 754 | 499.2 | 1.51 |
| M13 | 26 | 19 | 73% | 21 | 81% | 34 | 497.1 | 0.07 |
| M20 | 5 | 4 | 80% | 4 | 80% | 5 | 497.1 | 0.010 |
| M12 | 11 | 10 | 91% | 11 | 100% | 1,479 | 498.0 | 2.97 |

| | | | | | | | | |
|---|---|---|---|---|---|---|---|---|
| All 21 animals | 221 | 141 | 64% | 195 | 88% | 5,544 | 10,461 | 0.53 |

Supplementary Table 3. Detectors trained with each 60-s window standardized on its own (per-window scaling), compared with the per-recording detector of the main text. The control was trained on the same annotations with the same code and split as the per-recording detector and differs only in scaling. The production detector is the earlier released model, trained on the same annotations with an earlier version of the training code and a different validation split (1,561 against 1,451 validation windows). All three at detection threshold 0.5 and boundary threshold 0.1. The matched-count row keeps each detector's 2,499 most confident detections, the production detector's count, as an approximation of a higher threshold (boundaries are not recomputed). Not every held-out control detection was reviewed. Source detectors only.

| | Production per-window detector (earlier code) | Per-window control (same code) | Per-recording detector (main text) |
|---|---|---|---|
| Reference seizures detected at 0.5 / 0.1 (of 221) | 34 (15%) | 189 (86%) | 141 (64%) |
| Detections (per recording-hour) | 2,499 (0.24) | 29,258 (2.80) | 5,544 (0.53) |
| Recording time flagged | 13.2 h | 73.9 h | 17.6 h |
| Reference seizures among the 2,499 most confident detections | 34 | 37 | 88 |
| Animals with no detected reference seizure (of 11) | 4 | 0 | 0 |
| Held-out reference seizures detected (of 30) | 3 | 24 | 19 |
| Held-out detections that were seizures | 35/64 (55%) | 110 of 153 reviewed (72%), 378 not reviewed | 90/125 (72%) |

Supplementary Table 4. Held-out recordings per animal: reference seizures detected by each detector, the retrained detector's detections, and confirmed seizures after reviewing every detection. Data of Figs. 4B and 4C.

| **Animal** | **Reference seizures on held-out recordings** | **Source detected** | **Retrained detected** | **Retrained detections** | **Of which seizures** | **Precision** |
|---|---|---|---|---|---|---|
| M01 | 0 | – | – | 15 | 15 | 100% |
| M03 | 10 | 7 | 8 | 46 | 38 | 83% |
| M04 | 2 | 1 | 2 | 2 | 2 | 100% |
| M05 | 10 | 3 | 9 | 37 | 29 | 78% |
| M09 | 0 | – | – | 7 | 2 | 29% |
| M12 | 0 | – | – | 40 | 29 | 73% |
| M13 | 4 | 4 | 4 | 5 | 5 | 100% |
| M14 | 0 | – | – | 2 | 2 | 100% |
| M15 | 0 | – | – | 1 | 1 | 100% |
| M17 | 0 | – | – | 1 | 1 | 100% |
| M19 | 2 | 2 | 2 | 3 | 2 | 67% |
| M20 | 2 | 2 | 2 | 5 | 5 | 100% |
| M21 | 0 | – | – | 29 | 11 | 38% |
| All | 30 | 19 | 27 | 193 | 142 | 74% |

Supplementary Table 5. The convulsive classifier (Stage 2) applied to the retrained detector's detections, by Stage 2 threshold. Left, all second-cohort recordings: reference seizures kept and precision against the reference (among detections overlapping an adjudicated candidate, the share that are reference convulsive seizures). Right, held-out recordings: detections passing, confirmed seizures, convulsive seizures (overlapping a reference seizure or flagged convulsive by the reviewer), and reference seizures kept. Data of Fig. 7.

| **Stage 2 threshold** | **Reference seizures kept (of 221)** | **Precision vs reference** | **Held-out detections** | **Seizures** | **Convulsive** | **Held-out reference seizures kept (of 30)** |
|---|---|---|---|---|---|---|
| none | 195 (88%) | 9% | 193 | 142 (74%) | 37 (19%) | 27 |
| 0.3 | 187 (85%) | 10% | 145 | 112 (77%) | 37 (26%) | 27 |
| 0.45 | 164 (74%) | 12% | 117 | 98 (84%) | 34 (29%) | 25 |
| 0.55 | 152 (69%) | 16% | 87 | 74 (85%) | 30 (34%) | 24 |
| 0.65 | 133 (60%) | 24% | 58 | 53 (91%) | 27 (47%) | 22 |
| 0.75 | 91 (41%) | 32% | 40 | 40 (100%) | 23 (58%) | 19 |
| 0.8 (reported) | 70 (32%) | 39% | 26 | 26 (100%) | 19 (73%) | 17 |
| 0.9 | 28 (13%) | 60% | 14 | 14 (100%) | 11 (79%) | 10 |
| 0.95 | 18 (8%) | 82% | 7 | 7 (100%) | 7 (100%) | 7 |

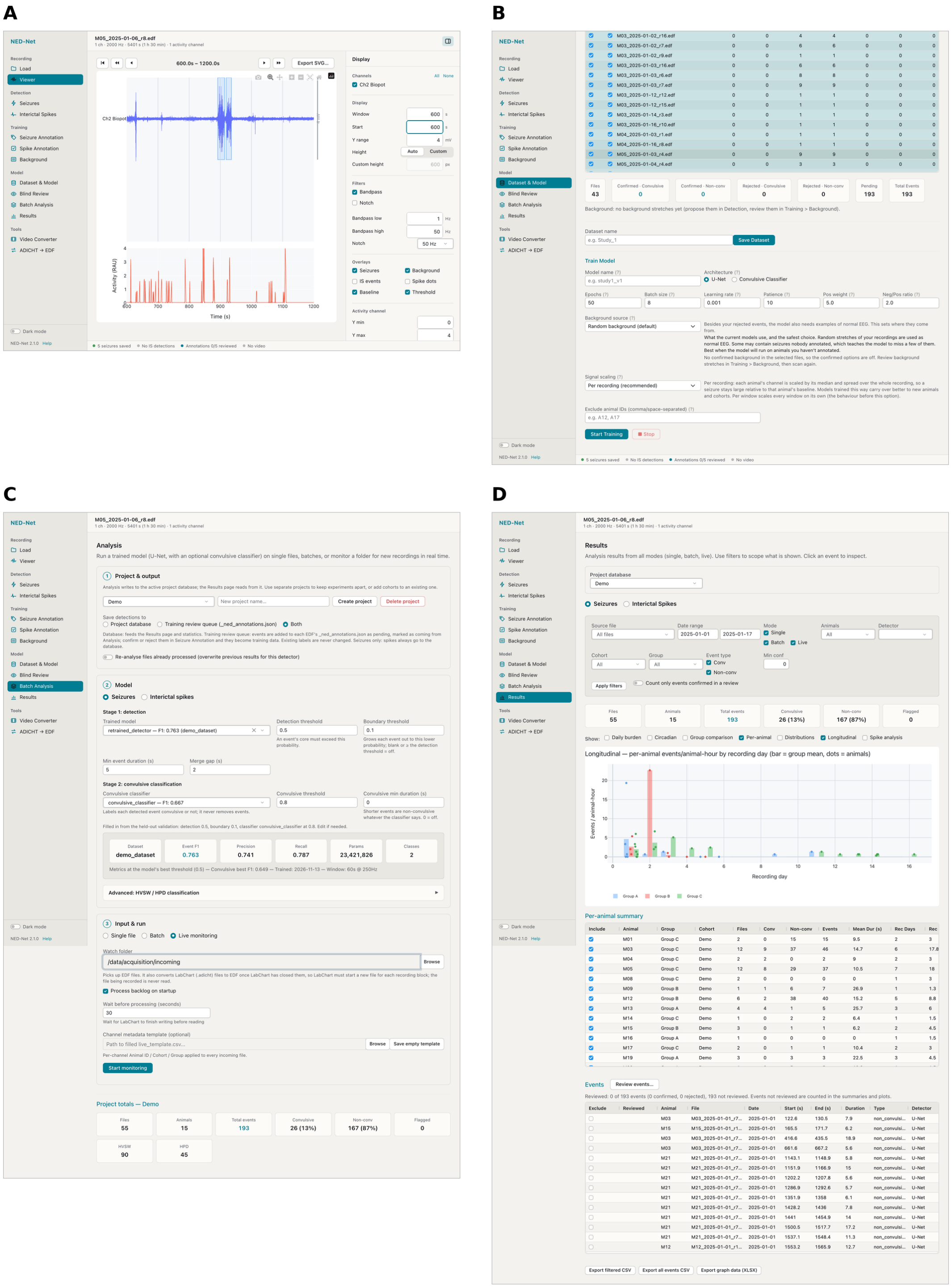


Supplementary Figure 1. Full views of four NED-Net pages, from NED-Net 2.1.0 with a demonstration project. (A) Viewer: a 600-s window of one channel with band-pass and notch filters, overlays of seizures, background stretches, baseline and threshold, and the activity channel below. (B) Dataset & Model: the per-file table of confirmed, rejected and pending events in the selected recordings, dataset export, and the training form with architecture, hyperparameters, background source and signal scaling. (C) Analysis: project and output selection, the Stage 1 detector and Stage 2 classifier with their operating point, the input mode with the live-monitoring settings, and the project totals. (D) Results: filters by file, date, mode, animal, cohort, group and event type, summary tiles, the view selector, the longitudinal graph, the per-animal summary table, the event list with the review control, and export of the filtered events and of the graph data.
